# Precision DNA Impurity Reduction Approaches for Ultra-Pure rAAV Manufacturing

**DOI:** 10.64898/2026.04.07.716878

**Authors:** Jingluan Han, Huan Chen, Xinping Tan, Zhiyong Dai, Ye Bu, HuaPeng Li

## Abstract

Recombinant adeno-associated virus (rAAV) vectors are a leading platform for gene delivery in basic and clinical research, yet large-scale manufacturing remains constrained by residual nucleic-acid impurities that compromise safety. In this study, we profiled the DNA species packaged within rAAV capsids and identified plasmid backbone sequences and host cell genomic DNA (hcDNA) as predominant contaminants. To mitigate this critical quality attribute, we implemented upstream strategies designed to fragment or excise backbone DNA, including TelN/TelROL excision, I-SceI meganuclease digestion, CRISPR/Cas9 cleavage, and Cre/LoxP recombination. Quantitatively, TelN/TelROL and I-SceI reduced encapsidated plasmid backbone DNA to approximately 20–30% and 20–40% of baseline levels, respectively, while CRISPR/Cas9 lowered it to about 10–20%. Notably, the Cre/LoxP system eliminated detectable plasmid backbone DNA without compromising vector-genome titers, indicating preserved genomic integrity. Additionlly, supplementating cell culture with a caspase inhibitor significantly reduced hcDNA contamination in rAAV particles to 1-5% of the baseline level. Collectively, these interventions provide practical bioprocess frameworks that markedly enhance rAAV purity *via* targeted DNA minimization and prevention of hcDNA fragmentation, thereby strengthening the safety profile of rAAV therapeutics in alignment with current Good Manufacturing Practice (cGMP) expectations.

## INTRODUCTION

Recombinant adeno-associated virus (rAAV) has become a cornerstone for gene delivery in both fundamental research and clinical therapy, owing to its favorable safety profile and efficient transduction (Wang et al., 2019). The clinical impact of rAAV is underscored by approved therapies such as Luxturna for Leber congenital amaurosis, Zolgensma for spinal muscular atrophy, and Hemgenix for hemophilia B (Mendell et al., 2017; Russell et al., 2017; Maguire et al., 2019). As numerous preclinical and clinical programs continue to advance, there is a pressing need to improve rAAV manufacturing with respect to productivity, scalability, and product purity (Tai et al., 2018; Srivastava et al., 2021).

In mammalian systems—most commonly HEK293-derived cells—rAAV is typically produced by transient co-transfection with three plasmids: (i) a gene-transfer plasmid carrying the transgene cassette flanked by AAV inverted terminal repeats (ITRs), (ii) a packaging plasmid encoding rep and cap, and (iii) a helper plasmid supplying functions required for viral replication and assembly (Ferrari et al., 1997; Fu et al., 2023). Alternative platforms include stable producer cell lines, baculovirus/Sf9 expression systems, and, more recently, cell-free approaches (Clément & Grieger, 2016; Kotin & Snyder, 2017; Brookwell et al., 2021). Regardless of platform, a central challenge remains: safeguarding the genetic purity of the final rAAV product. Beyond the intended rAAV genomes carrying the transgene cassette, purified vector preparations can harbor unintended DNA species derived from production plasmids and host-cell genomes (hcDNA) (Wright et al., 2014; Lecomte et al., 2015; Tai et al., 2018; Tran et al., 2022; Brimble et al., 2023). These impurities raise safety concerns for clinical translation for several reasons (Wright et al., 2014; Srivastava et al., 2021; Brimble et al., 2022): (i) residual DNA may be transferred *in vivo* and provoke immune responses to foreign sequences or encoded proteins; (ii) inadvertent expression of contaminant-derived genes could trigger off-target biological effects; and (iii) increased genomic heterogeneity can undermine vector performance, rasing variability in therapeutic outcomes.

Accordingly, there is a pressing need for improved manufacturing strategies that minimize encapsidated DNA impurities in rAAV products. Deep sequencing of purified rAAV lots has clarified both the composition and burden of encapsidated off-target DNA. Multiple studies identify sequence-specific drivers of mispackaging—most notably the inverted terminal repeats (ITRs) and the P5 promoter—as recurrent hotspots (Guerin et al., 2020; Radukic et al., 2020; Tran et al., 2022; Brimble et al., 2022; Taylor et al., 2024). The ITRs are the essential *cis*-acting elements that direct genome replication and packaging, while the P5 promoter controlling Rep expression contains a Rep-binding element (RBE) that is structurally homologous to the RBE within the ITRs (Kotin, 1994; McCarty, 2008). This shared recognition architecture is thought to aberrantly recruit the replication/packaging machinery to non-transgene sequences, thereby facilitating inappropriate encapsidation, although the precise mechanisms remain under active investigation.

Multiple plasmid-engineering strategies have been explored to curb unwanted DNA in purified rAAV. Adding a spacer adjacent to the ITRs or the P5 promoter can, in some contexts, diminish plasmid-backbone encapsidation by increasing the separation between these packaging-competent elements and the backbone sequence (Brimble et al., 2022; Taylor et al., 2024). Likewise, enlarging the nonessential backbone with a “stuffer” insert reduces its recovery in final products by exploiting rAAV’s ∼4.7-kb packaging constraint (Taylor et al., 2024). However, these interventions are not uniformly effective and may trade off with vector yield or introduce new impurities (Taylor et al., 2024).

A more radical line of work aims to remove the plasmid backbone entirely. Minicircle DNA—generated in *E. coli via* L-arabinose–induced, ParA resolvase–mediated site-specific recombination—excises bacterial elements before transfection, yielding a circular construct that retains only the sequences required for rAAV production and markedly lowers backbone mispackaging (Schnödt et al., 2016). In parallel, linear, covalently closed–ended DNA (dbDNA) produced *in vitro* with Φ29 DNA polymerase and a protelomerase provides a backbone-free substrate that similarly prevents backbone encapsidation, while offering a fundamentally different manufacturing paradigm that bypasses conventional plasmid preparation (Karbowniczek et al., 2017). Both platforms have shown substantial reductions in contaminant DNA and are emerging as next-generation inputs for safer, more efficient vector manufacture. Nonetheless, producing these alternative DNA forms typically necessitates additional purification and handling steps during rAAV preparation, introducing process complexity and potential cost trade-offs.

Here, we develop and optimize upstream strategies designed to generate miniDNA and preventing hcDNA fragmentation during rAAV manufacture, with the goal of further reducing residual DNA contaminants. By engineering plasmid architecture, incorporating emerging DNA substrates, and systematically benchmarking their effects on vector purity, we outline a framework for robust, scalable, and clinically acceptable rAAV production. Adoption of these refinements will be critical to improving the safety and lot-to-lot consistency of gene therapies as they advance toward broader clinical use.

## Results

### Plasmid-Backbone and Host cell DNA–Derived Contaminants in rAAV Preparations

To characterize DNA species packaged in rAAV particles, we used next-generation sequencing (NGS) to quantify encapsidated sequences. rAAV-eGFP was produced by co-transfecting HEK293 cells with pGOI-eGFP/pRepCap/pHelper; empty particles (rAAV-Em) were generated with pRepCap/pHelper. DNA extracted from purified particles was sequenced, and reads were aligned to the three production plasmids and the human reference genome.

In rAAV-eGFP, 98.732% of reads mapped to the ITR-to-ITR region of pGOI-eGFP (pGOI-eGFP_Specific). Reads mapping to plasmid backbones totaled 0.675%, including 0.337% to the RepCap region (pRepCap_Specific) and 0.002% to the pHelper-specific region; 0.548% aligned to hcDNA (Table 1). In rAAV-Em, 75.828% of reads mapped to the pRepCap-specific region, 21.203% to pRepCap/pHelper backbones, 0.015% to the pHelper-specific region, and 3.473% to the host genome (Table 1).

**Table 1.** Distribution of NGS reads aligned to production plasmids and host genome in purified rAAV preparations.

| Feature | rAAV-eGFP | rAAV-Em |
| --- | --- | --- |
| pGOI-eGFP_Specific | 98.732% | / |
| pRepCap_Specific | 0.337% | 75.828% |
| pHelper_Specific | 0.002% | 0.015% |
| Plasmid backbone | 0.675% | 21.203% |
| Host cell DNA | 0.548% | 3.473% |

We next performed quantitative PCR (qPCR) to assess specific plasmid components in rAAV-eGFP and rAAV-Em, using equal particle inputs (1.0×10^8 capsids). Primer pairs targeted WPRE (pGOI-eGFP specific), ori (backbone common to pGOI-eGFP/pRepCap/pHelper), Rep and Cap (pRepCap specific), and E2A/E4/VARNA (pHelper specific) (Figure S1A). In rAAV-eGFP, WPRE was most abundant, followed by ori, Cap, and Rep. In rAAV-Em, Cap and Rep exceeded ori. E2A, E4, and VARNA were undetectable in both particle types (Figure S1B). Together, these data indicate that the principal contaminants arise from plasmid backbones and hcDNA. Consistent with prior reports, we hypothesize that residual plasmid DNA is preferentially co-packaged *via* interactions involving the AAV ITRs (from pGOI-eGFP) and/or the P5 promoter in pRepCap.

### Reducing Residual Plasmid DNA in rAAV Production with the TelN/TelROL System

To diminish plasmid-backbone encapsidation, we applied a TelN/TelROL excision strategy to generate miniDNA during rAAV manufacture. First, two TelROL recognition sites were placed immediately upstream of the 5′ ITR and downstream of the 3′ ITR in pGOI-LucGFP to create pGOI-TelROL. The plasmid was digested *in vitro* with TelN protelomerase and co-transfected with pRepCap and pHelper. Relative to the undigested control (ND), TelN digestion (+TelN) reduced the WPRE titer (ITR-to-ITR genome proxy) and ori titer (backbone proxy) to 0.55× and 0.13×, respectively, lowering the ori/WPRE ratio to 0.23× (Figure 1A). Thus, *in vitro* TelN processing decreased residual backbone DNA, albeit with a concomitant drop in vector-genome titer.

**Figure 1.**
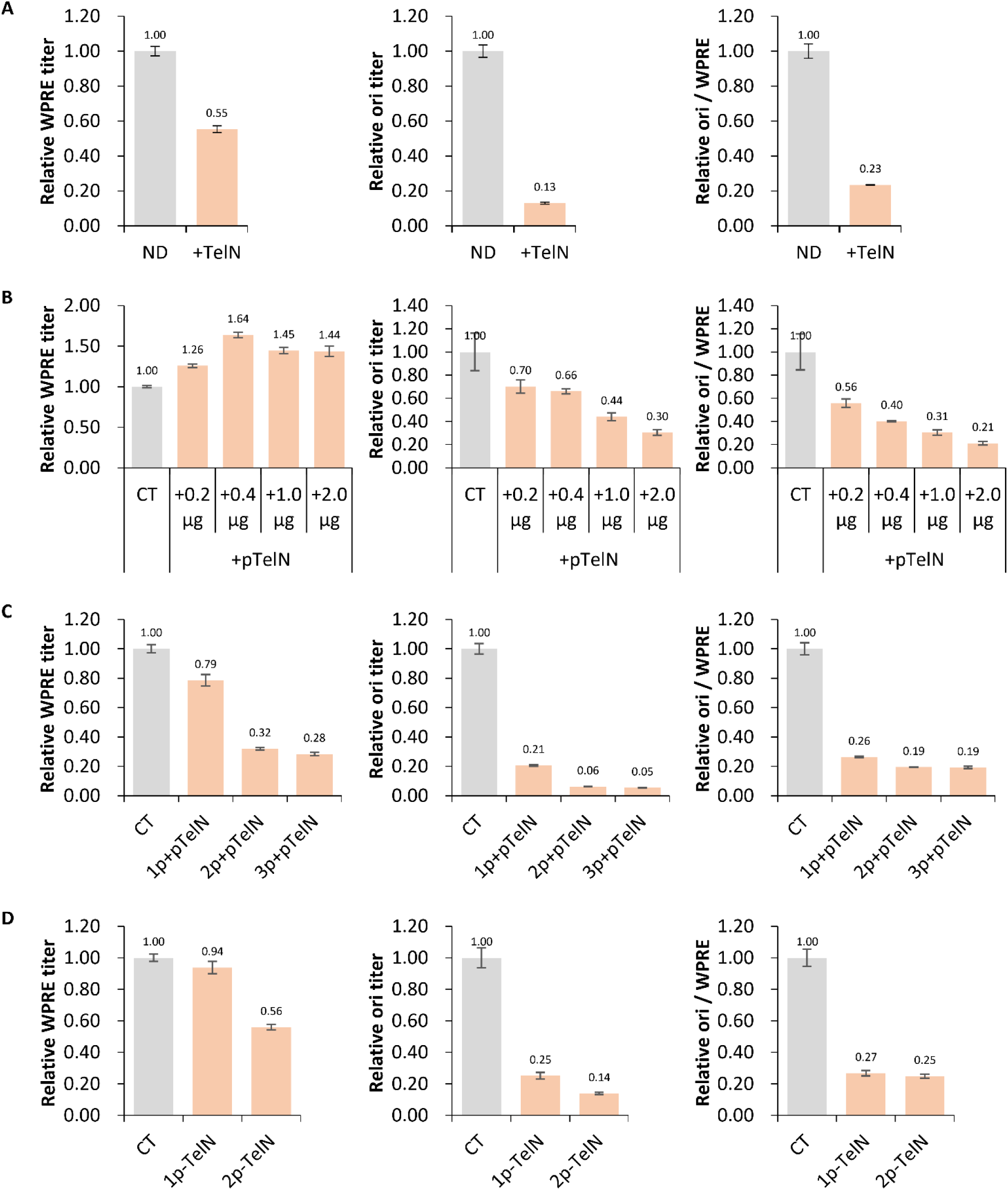
TelN/TelROL excision strategy reduces residual plasmid DNA during rAAV production. (A) rAAV was produced by co-transfecting pRepCap and pHelper with either TelN-digested pGOI-TelROL (+TelN) or undigested pGOI-TelROL (ND). (B) Dose–response of TelN expression: increasing amounts of pTelN (+pTelN) were added to the standard three-plasmid system (pGOI-TelROL/pRepCap/pHelper); CT denotes the three-plasmid co-transfection without pTelN. (C) Aplication of TelROL sites to one, two, or three plasmids: rAAV was produced with pGOI-TelROL, pRepCap-TelROL, pHelper-TelROL, and pTelN. Conditions were: CT (pGOI-TelROL/pRepCap/pHelper), 1p+pTelN (pGOI-TelROL/pRepCap/pHelper/pTelN), 2p+pTelN (pGOI-TelROL/pRepCap-TelROL/pHelper/pTelN), and 3p+pTelN (pGOI-TelROL/pRepCap-TelROL/pHelper-TelROL/pTelN). (D) TelN expression cassette engineered on the helper plasmid: rAAV was produced using pGOI-TelROL, pRepCap-TelROL, and pHelper-TelN. Conditions were: CT (pGOI-TelROL/pRepCap/pHelper), 1p-TelN (pGOI-TelROL/pRepCap/pHelper-TelN), and 2p-TelN (pGOI-TelROL/pRepCap-TelROL/pHelper-TelN). For each condition, genome titer (WPRE) and plasmid-backbone titer (ori) were quantified by qPCR. Relative WPRE titer, relative ori titer, and the ori/WPRE ratio were calculated against the corresponding control rAAV samples.

We next expressed TelN *in vivo*. Introducing a TelN expression plasmid (pTelN) during packaging increased WPRE titers (1.26–1.64×) while decreasing ori titers (0.70–0.30×), yielding a dose-dependent reduction in ori/WPRE (Figure 1B). This indicates that intracellular TelN excision more effectively curbs backbone carryover while sustaining (and in some conditions improving) genome output.

To localize sources of residual DNA, we inserted TelROL sites into additional plasmids: pRepCap-TelROL (TelROL upstream of Rep and downstream of P5) and pHelper-TelROL (TelROL upstream of E2A and downstream of VARNA). Compared with control (CT), co-transfection of pGOI-TelROL/pRepCap/pHelper/pTelN (1p + pTelN) yielded 0.79× WPRE, 0.21× ori, and 0.26× ori/WPRE; co-transfection of pGOI-TelROL/pRepCap-TelROL/pHelper/pTelN (2p + pTelN) yielded 0.32× WPRE, 0.06× ori, and 0.19× ori/WPRE; co-transfection of pGOI-TelROL/pRepCap-TelROL/pHelper-TelROL/pTelN (3p + pTelN) yielded 0.28× WPRE, 0.05× ori, and 0.19× ori/WPRE (Figure 1C).

These results suggest that separating the backbone from pGOI substantially reduces residual plasmid DNA with only a modest impact on genome titer, whereas separation in pRepCap further suppresses contaminants but more strongly depresses genome yield. Modifying pHelper did not meaningfully reduce backbone carryover. Finally, we embedded the TelN cassette in pHelper (pHelper-TelN) to streamline expression. Relative to CT, 1p-TelN (pGOI-TelROL/pRepCap/pHelper-TelN) produced 0.94× WPRE and 0.25× ori (ori/WPRE = 0.27), while 2p-TelN (pGOI-TelROL/pRepCap-TelROL/pHelper-TelN) yielded 0.56× WPRE and 0.14× ori (ori/WPRE = 0.25) (Figure 1D). These data corroborate that excising backbones from pGOI and/or pRepCap is the principal lever for reducing residual plasmid DNA.

In conclusion, the TelN/TelROL approach generates miniDNA and effectively lowers plasmid-backbone contamination in rAAV preparations. Residual DNA predominantly originates from pGOI, followed by pRepCap, with minimal contribution from pHelper.

### Reducing Plasmid Contaminants in rAAV Production with the I-SceI Meganuclease

We evaluated I-SceI and its cognate recognition site to cleave production plasmids during rAAV manufacture. Two I-SceI sites were inserted immediately upstream of the 5′ ITR and downstream of the 3′ ITR in pGOI-LucGFP to generate pGOI-IRS. Following *in vitro* digestion with I-SceI, pGOI-IRS was co-transfected with pRepCap and pHelper. Relative to the undigested control (ND), the WPRE titer (ITR-to-ITR genome proxy) and ori titer (backbone proxy) decreased to 0.75× and 0.17×, respectively, yielding an ori/WPRE ratio of 0.23× (Figure 2A), indicating reduced backbone carryover.

**Figure 2.**
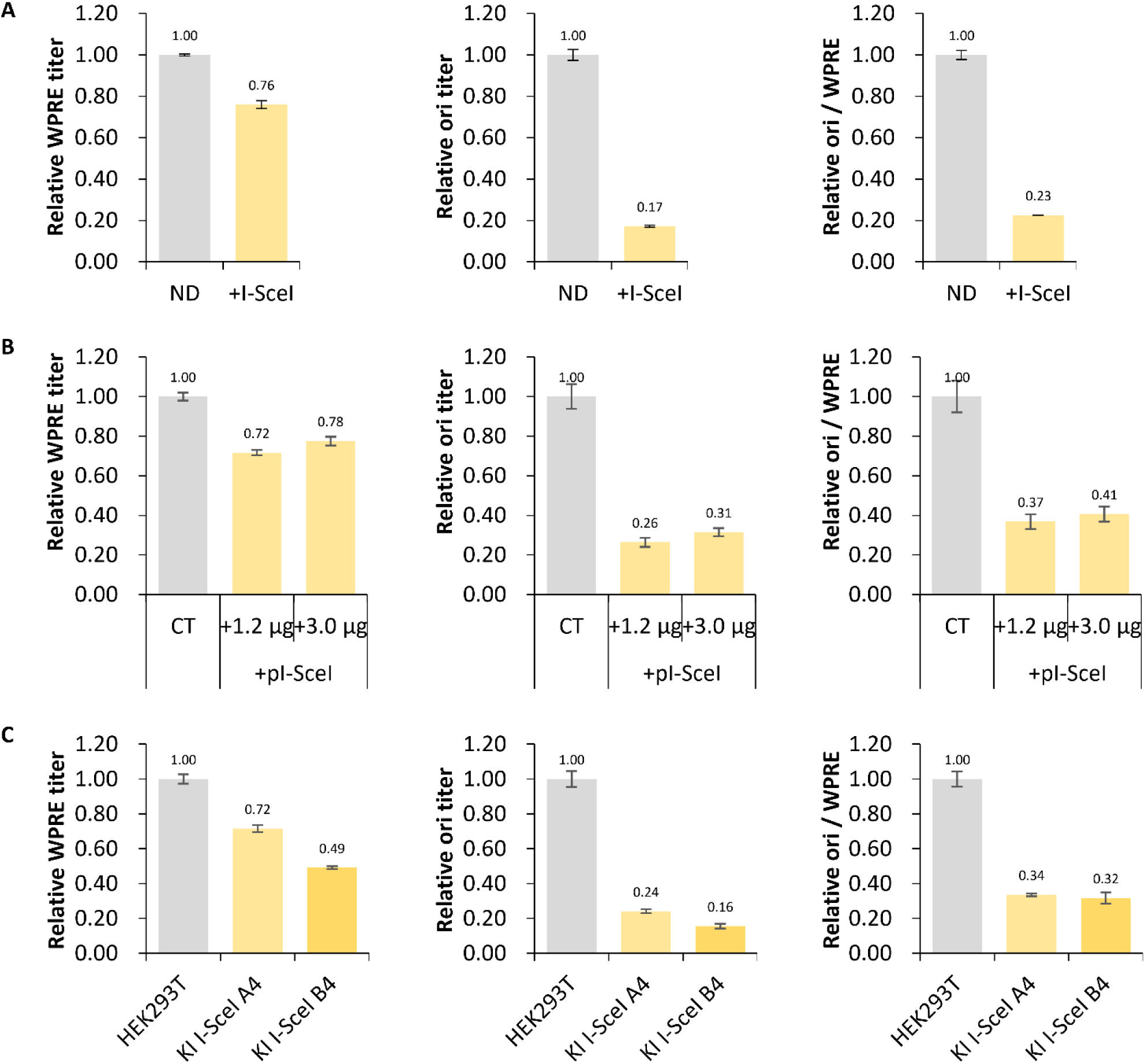
I-SceI meganuclease digestion reduces plasmid-derived contaminants during rAAV production. (A) rAAV was produced by co-transfecting pRepCap and pHelper with either I-SceI–digested pGOI-IRS (+I-SceI) or undigested pGOI-IRS (ND). (B) Dose–response of I-SceI expression: increasing amounts of pI-SceI (+pI-SceI) were added to the standard three-plasmid system (pGOI-IRS/pRepCap/pHelper); CT denotes the three-plasmid co-transfection without pI-SceI. (C) rAAV production in KI-I-SceI cells versus wild-type HEK293T controls using pGOI-IRS, pRepCap, and pHelper. For each condition, genome titer (WPRE) and plasmid-backbone titer (ori) were quantified by qPCR. Relative WPRE titer, relative ori titer, and the ori/WPRE ratio were calculated against the corresponding control rAAV samples.

To enable *in vivo* cleavage, we introduced an I-SceI expression plasmid (pI-SceI) during packaging. Compared with control (CT), WPRE decreased to 0.72–0.78×, ori to 0.26–0.31×, and ori/WPRE to 0.37–0.41× (Figure 2B). We further generated HEK293T knock-in cell lines (KI I-SceI A4 and B4) stably expressing I-SceI. Using these cells, WPRE fell to 0.72× and 0.49×, ori to 0.24× and 0.16×, and ori/WPRE to 0.34× and 0.32×, respectively (Figure 2C).

These data suggested that I-SceI–mediated cleavage of production plasmids consistently lowers plasmid-backbone contamination, with modest impacts on vector-genome titers.

### Lowering DNA Impurities in rAAV Production with CRISPR/Cas9

We evaluated the CRISPR/Cas9 system as a means to degrade residual plasmid DNA during rAAV manufacture. Two identical target sites (Tgfp) were inserted immediately upstream of the 5′ ITR and downstream of the 3′ ITR in pGOI-mChe, and a corresponding sgRNA was encoded on the plasmid backbone to generate pGOI-Tgfp. A pCas9 plasmid was used to supply Cas9. Co-transfection of pCas9 with pGOI-Tgfp/pRepCap/pHelper increased the WPRE titer (3.60× and 3.27× vs. control, CT) and reduced ori (backbone proxy) to 0.35× and 0.53×, lowering the ori/WPRE ratio to 0.10× and 0.16× (Figure 3A). Embedding the Cas9 cassette in pHelper (pHelper-Cas9) and co-transfecting pGOI-Tgfp/pRepCap/pHelper-Cas9 decreased WPRE and ori to 0.35× and 0.05×, respectively (ori/WPRE = 0.14×) relative to CT (Figure 3B), indicating on-process backbone suppression with a trade-off in genome output.

**Figure 3.**
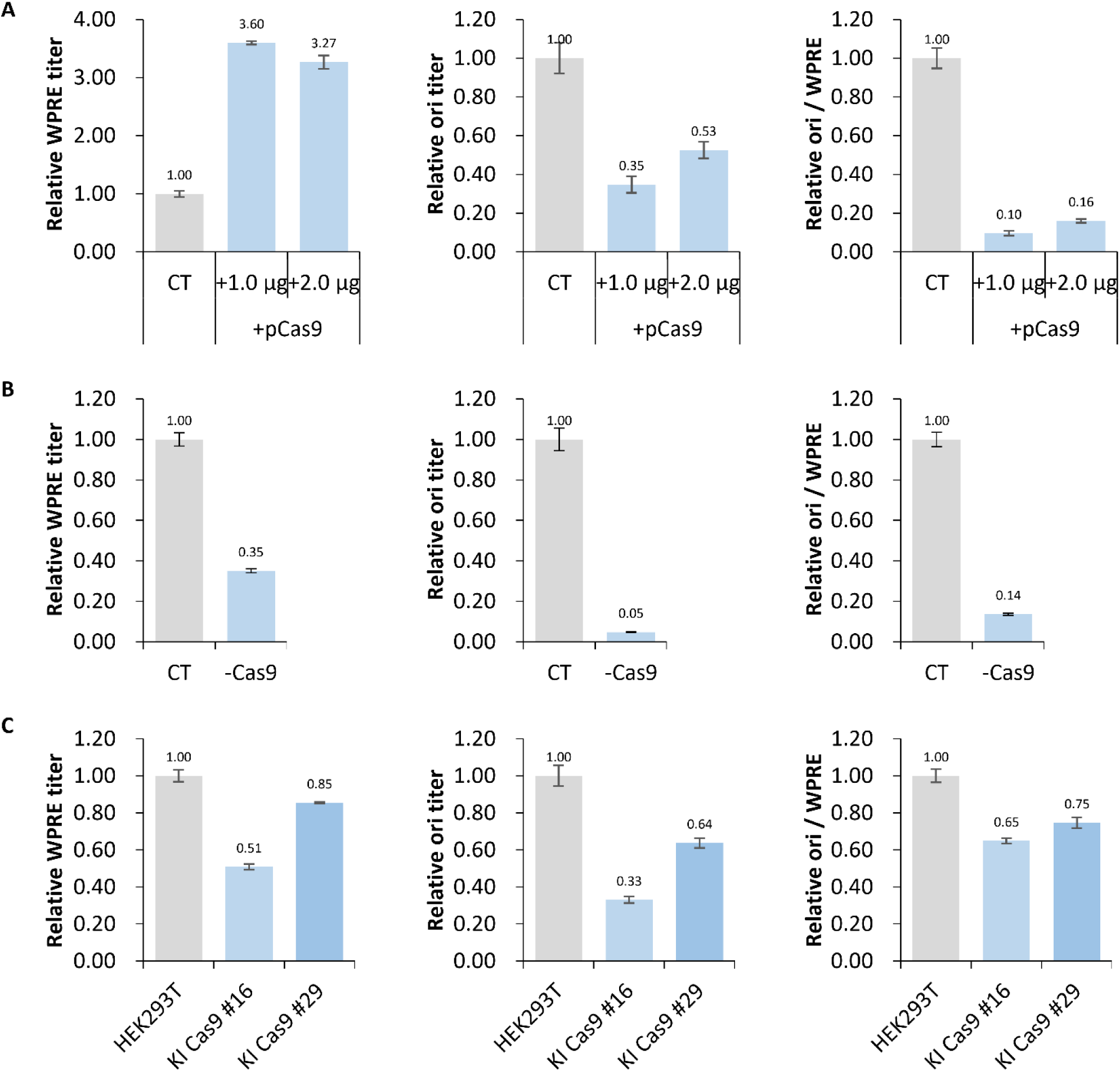
CRISPR/Cas9-mediated cleavage reduces DNA impurities during rAAV production. (A) rAAV was produced by adding the pCas9 plasmid (+pCas9) to the standard three-plasmid system (pGOI-Tgfp/pRepCap/pHelper); CT denotes the three-plasmid co-transfection without pCas9. (B) rAAV was generated using pGOI-Tgfp, pRepCap, and a helper plasmid bearing Cas9 (pHelper-Cas9). The "−Cas9" condition indicates absence of a separate pCas9 plasmid); CT represents pGOI-Tgfp/pRepCap/pHelper. (C) rAAV packaging in Cas9 knock-in (KI-Cas9) cells versus wild-type HEK293T controls using pGOI-Tgfp, pRepCap, and pHelper. For all conditions, genome titer (WPRE) and plasmid-backbone titer (ori) were quantified by qPCR. Relative WPRE, relative ori, and the ori/WPRE ratio were calculated against the corresponding control rAAV samples.

To test constitutive nuclease expression, we generated HEK293T Cas9 knock-in lines (KI Cas9 #16 and #19) (Figure S3A–S3D). Functional activity was confirmed by reduced GFP fluorescence after pGFP + sgGFP transfection and by T7E1 cleavage (Figure S3E–S3F). However, using these KI lines for rAAV production with pGOI-Tgfp/pRepCap/pHelper yielded only modest changes *vs.* parental HEK293T: WPRE 0.51×/0.85×, ori 0.33×/0.64×, and ori/WPRE 0.65×/0.75× (Figure 3C).

While transient or helper-encoded Cas9 can reduce backbone carryover, constitutive Cas9 expression confers only limited benefit. Overall, CRISPR/Cas9 is not an optimal standalone strategy for minimizing plasmid DNA contamination in rAAV manufacturing.

### Improving rAAV DNA Purity with the Cre/LoxP System

To leverage the relative stability of circular DNA over linear DNA, we used the Cre/LoxP system to fragment production plasmids into two minicircles during rAAV manufacturing. Two loxP sites were placed upstream of the 5′ ITR and downstream of the 3′ ITR in pGOI-LucGFP to generate pGOI-loxP. Pre-incubation of pGOI-loxP with Cre recombinase increased the WPRE (ITR-to-ITR) titer to 1.56× of the non-recombined control (NR) and lowered the ori (backbone proxy) titer to 0.62×, reducing the ori/WPRE ratio to 0.40× (Figure 4A). Thus, *in vitro* Cre recombination modestly decreases residual plasmid DNA.

**Figure 4.**
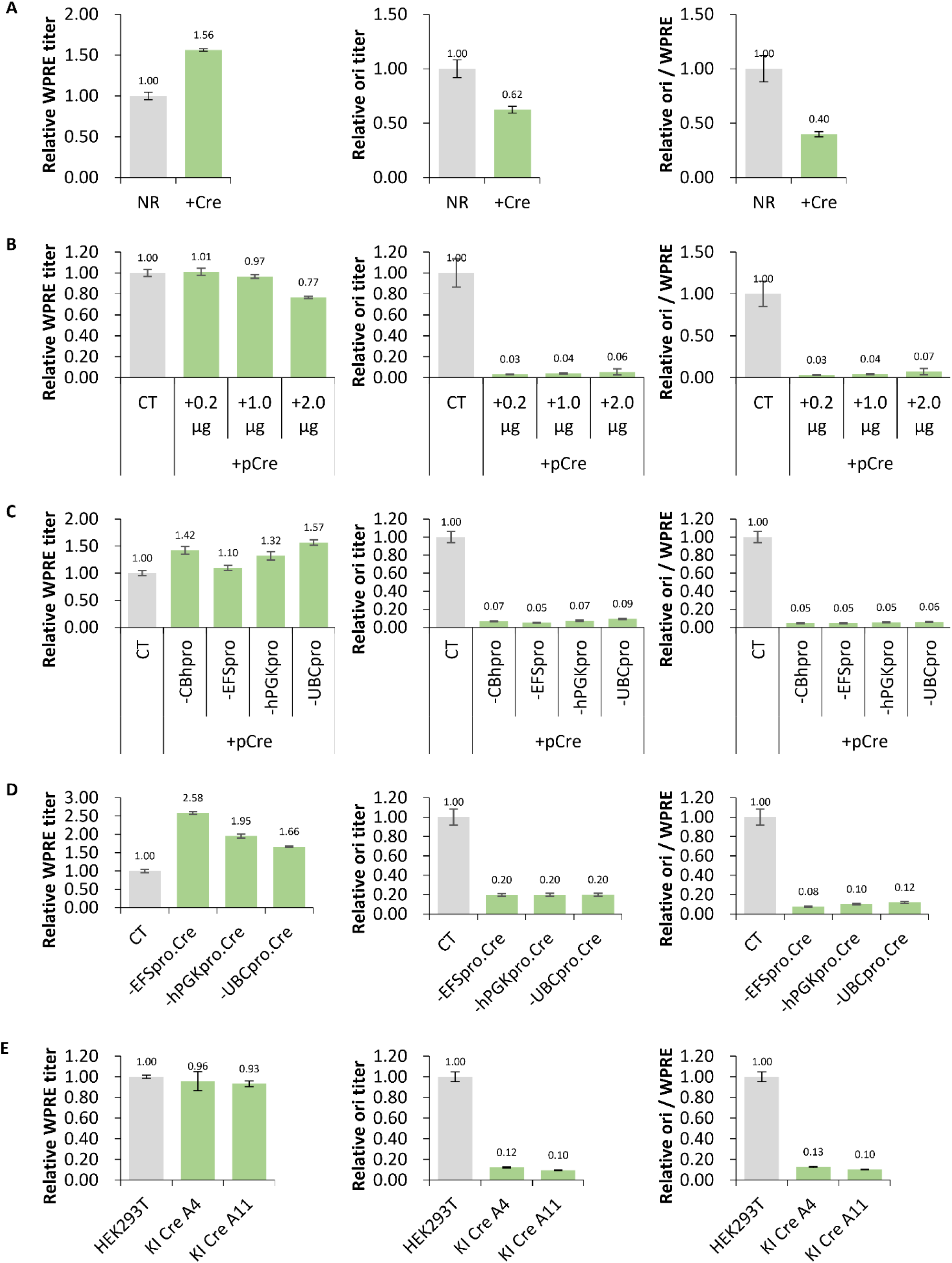
Improved rAAV DNA purity using the Cre/loxP system. (A) rAAV was produced by co-transfecting pRepCap and pHelper with either Cre-recombined pGOI-loxP (+Cre) or non-recombined pGOI-loxP (NR). (B) Titration of Cre expression: increasing amounts of pCre (+pCre) was added to the standard three-plasmid system (pGOI-loxP/pRepCap/pHelper); CT denotes the three-plasmid co-transfection without pCre. (C) The three-plasmid system (pGOI-loxP/pRepCap/pHelper) was supplemented with pCre constructs driven by different promoters; CT indicates pGOI-loxP/pRepCap/pHelper alone. (D) rAAV was produced using pGOI-loxP, pRepCap, and pHelper-Cre plasmids encoding Cre under control of various promoters. (E) rAAV production in Cre knock-in (KI-Cre) cells was compared with wild-type HEK293T controls using pGOI-loxP, pRepCap, and pHelper. For each condition, genome titer (WPRE) and plasmid-backbone titer (ori) were quantified by qPCR. Relative WPRE, relative ori, and the ori/WPRE ratio were calculated against the corresponding control rAAV samples.

For *in vivo* recombination, we expressed Cre from a CBh promoter (pCBhpro.Cre). Increasing pCre input produced a graded reduction in WPRE (1.01× → 0.97× → 0.77× of control) and a marked decrease in ori (0.06–0.03×), driving ori/WPRE as low as 0.03× (Figure 4B). This indicates that intracellular Cre efficiently suppresses backbone carryover, with lower Cre doses better preserving vector yield.

We next compared promoters driving Cre on a separate plasmid—CBh (strong), EFS and hPGK (medium), and UBC (weak). Co-transfection with these pCre variants increased WPRE (1.42×, 1.10×, 1.32×, and 1.57×, respectively) and reduced ori (0.07×, 0.05×, 0.07×, 0.09×), yielding ori/WPRE ≈ 0.05× across conditions (Figure 4C). These data reinforce that a controlled Cre level is optimal for minimizing backbone without sacrificing output. To streamline delivery, we embedded Cre cassettes into pHelper (EFS, hPGK, or UBC promoters), generating pHelper-EFSpro.Cre, pHelper-hPGKpro.Cre, and pHelper-UBCpro.Cre. Relative to control co-transfection (pGOI-loxP/pRepCap/pHelper), these helper-encoded constructs increased WPRE to 2.58×, 1.95×, and 1.66×, respectively, while reducing ori to 0.20× and ori/WPRE to 0.08, 0.10, and 0.12 (Figure 4D), confirming efficient on-process cleanup with modest promoter strength.

Finally, we generated Cre knock-in HEK293T lines (KI Cre A4 and A11). Transient pGOI-loxP transfection verified robust recombination, producing two smaller circles (Figure S4D–S4E). During rAAV production, these KI lines showed a slight WPRE decrease (∼0.9×) but a pronounced reduction in ori (0.12× and 0.10×) and ori/WPRE (0.13× and 0.10×) versus parental cells (Figure 4E).

The Cre/LoxP strategy reliably converts production plasmids into minicircles, substantially lowering plasmid-backbone contamination while preserving or improving vector-genome output when Cre expression is properly tuned. Embedding Cre on the helper plasmid or using Cre-expressing producer cells provides practical routes for consistent, high-purity rAAV manufacture.

### NGS Profiling of DNA Impurities in rAAV Produced *via* the Cre/LoxP System

To assess how Cre/LoxP impacts encapsidated DNA contaminants, we produced three rAAV preparations in HEK293 cells: CT (pGOI-loxP/pRepCap/pHelper), 1p+Cre (pGOI-loxP/pRepCap/pHelper-Cre), and 2p+Cre (pGOI-loxP/pRepCap-loxP/pHelper-Cre). Relative to CT, 1p+Cre increased the WPRE titer to 1.11×, whereas 2p+Cre reduced it to 0.48×; ori (backbone proxy) fell sharply to 0.06× and 0.01×, respectively, lowering ori/WPRE to 0.05× (1p+Cre) and 0.02× (2p+Cre) (Figure S5).

Next-generation sequencing of purified particles, with reads mapped to production plasmids and the human reference genome, showed that pGOI-eGFP_Specific (ITR-to-ITR) content rose slightly in 1p+Cre (97.683%) and 2p+Cre (99.168%) versus CT (97.125%). In contrast, plasmid-backbone representation dropped from 0.186% (CT) to 0.006% (1p+Cre) and 0.004% (2p+Cre) (Table 2).

**Table 2.** Proportion of NGS reads in rAAV particles produced using the Cre/loxP system.

| Feature | rAAV-CT | rAAV-1p+Cre | rAAV-2p+Cre |
| --- | --- | --- | --- |
| pGOI-eGFP_Specific | 97.125% | 97.683% | 99.168% |
| pRepCap_Specific | 0.117% | 0.147% | 0.139% |
| pHelper_Specific | 0.002% | 0.001% | 0.000% |
| Plasmid backbone | 0.186% | 0.006% | 0.004% |
| Host cell DNA | 2.664% | 2.285% | 0.799% |

Fragmenting production plasmids into two circular molecules *via* Cre/LoxP markedly reduces backbone carryover while maintaining—or in some contexts improving—on-target genome packaging, underscoring this approach as a practical route to higher-purity rAAV.

### Reduction of hcDNA in rAAV particles by Caspase Inhibitor Supplementation

To investigate the effect of small-molecule compounds on hcDNA packaging during rAAV production, we screened a panel of apoptosis inhibitors, focusing primarily on caspase inhibitors (Emricasan and Q-VD-OPh). These inhibitors were supplemented at varying concentrations into the cell culture medium throughout the rAAV transfection phase. The results showed that none of the caspase inhibitor treatments significantly affected the final genome titer of rAAV (Figure 5). However, substantial reductions in hcDNA levels were detected under specific conditions. Treatment with 3.0 μM and 5.0 μM Emricasan reduced residual hcDNA to 0.02- and 0.03-fold of the control level, whereas 2.0 μM and 5.0 μM Q-VD-OPh lowered hcDNA to 0.02- and 0.01-fold of the control (Figure 5).

**Figure 5.**
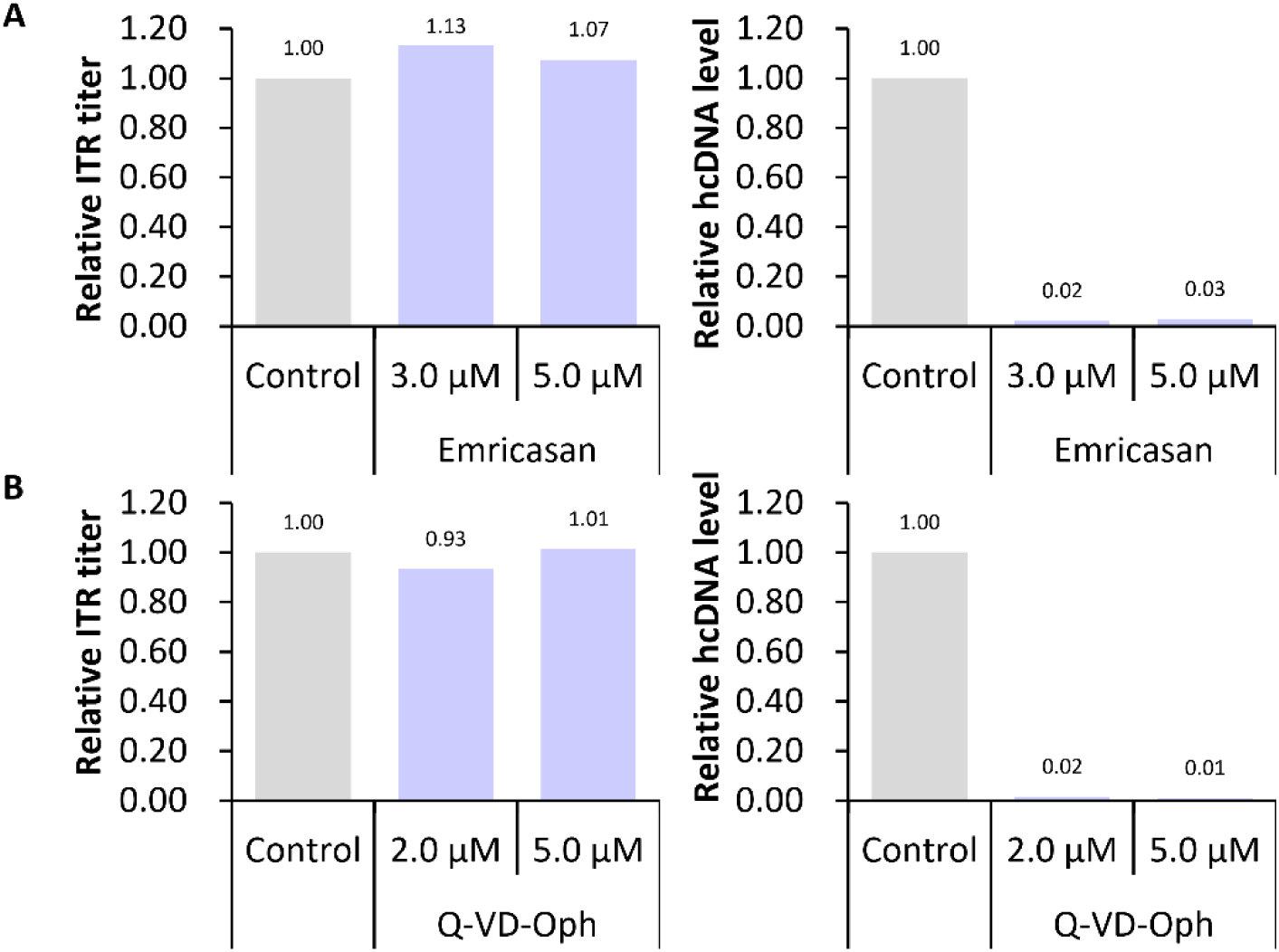
Reduction of hcDNA in rAAV particles by Caspase Inhibitor Supplementation. (A) Effect of Emricasan supplemention at indicated concentrations on rAAV genome titer and residual hcDNA levels. (B) Effect of Q-VD-OPh supplementation at indicated concentrations on rAAV genome titer and residual hcDNA levels.

Together, these findings demonstrate that selected caspase inhibitors can effectively decouple hcDNA NA packaging from viral genome production, providing a viable strategy to enhance the purity of rAAV preparations without compromising yield.

## Discussion

Residual DNA in rAAV preparations remains a major obstacle for gene therapy, with the potential to compromise both safety and efficacy. In this study, we systematically profiled encapsidated contaminants and benchmarked multiple upstream genetic-engineering strategies to mitigate this critical quality attribute. Our analyses show that impurities in purified rAAV are derived predominantly from plasmid backbones, Rep/Cap sequences, and hcDNA. Although on-target, ITR-to-ITR transgene reads were consistently abundant, backbone, RepCap, and hcDNA sequences were also detected (Tables 1–2), consistent with prior reports (Wright et al., 2014; Lecomte et al., 2015; Brimble et al., 2023). Notably, contaminant profiles differed between filled and empty particles: empty rAAV exhibited markedly higher proportions of pRepCap-specific reads (75.828%) and plasmid backbone (21.203%) (Table 1). Collectively, these findings support a model in which mispackaging is driven by *cis*-elements that recruit Rep—specifically, the AAV ITRs on the pGOI plasmid and the P5 promoter within pRepCap—aligning with prior evidence that ITR structures and P5-embedded RBE motifs promote unintended encapsidation *via* structural mimicry of AAV replication origins (Guerin et al., 2020; Radukic et al., 2020; Brimble et al., 2022).By deploying several strategies to physically separate the ITR-flanked rAAV genome from surrounding plasmid sequences—TelN/TelROL excision, I-SceI meganuclease digestion, CRISPR/Cas9 cleavage, and Cre/LoxP recombination—we achieved substantial reductions in plasmid-backbone carryover. The common principle of the first three approaches is site-specific cutting of the pGOI backbone to release the genome cassette and leave behind smaller, packaging-averse backbone fragments.

1. TelN/TelROL excision. Installing TelROL sites in pGOI (and, in some experiments, pRepCap) and supplying TelN lowered the ori/WPRE ratio to ∼0.2–0.3× of control with only a modest decrease in genome titer (Figure 1). In contrast, placing TelROL within pRepCap caused a pronounced loss of genome output (Figure 1C–D), underscoring the need to maintain intact Rep/Cap expression. Notably, separating the backbone from pHelper did not reduce residual contamination (Figure 1C), indicating that helper sequences contribute minimally to mispackaging and helping prioritize optimization efforts.
2. I-SceI meganuclease. I-SceI lowered ori/WPRE to ∼23–41% of control (Figure 2) with modest titer penalties. Although effective, its impact was smaller than other methods, likely reflecting incomplete cleavage and/or the reduced stability of linearized DNA and its greater susceptibility to degradation *in vivo*.
3. CRISPR/Cas9 cleavage. Cas9 expressed from a separate plasmid reduced ori/WPRE to 10–16% of control (Figure 3A), an effect retained when Cas9 was helper-encoded (∼14%; Figure 3B). However, constitutive Cas9 expression in knock-in lines produced only modest gains (65–75% of control; Figure 3C), implying that precise timing and dosing of Cas9 are critical. Potential off-target editing further limits the attractiveness of CRISPR/Cas9 as a standalone solution for clinical manufacturing.

The Cre/LoxP system proved the most effective approach for minimizing plasmid-backbone carryover. Flanking the pGOI ITRs with loxP sites and expressing Cre *in vivo* resolved the construct into two circular molecules—a minicircle bearing the GOI cassette and a separate backbone minicircle. *In vitro* Cre recombination reduced the ori/WPRE ratio to ∼40% of control (Figure 4A), whereas *in vivo* Cre lowered it to as little as 3% with minimal impact on vector-genome titers (Figure 4B). This benefit was robust across promoter strengths (CBh, EFS, hPGK, UBC; Figure 4C). Embedding Cre on the helper plasmid further improved performance, decreasing ori/WPRE to 8–12% of control while increasing WPRE titers 1.66–2.58× (Figure 4D)—suggesting that removing backbone sequences not only purifies the product but may also enhance packaging efficiency. Stable Cre producer lines yielded similar reductions (Figure 4E). Concordantly, NGS showed backbone reads falling from 0.186% (control) to 0.004–0.006% with Cre/LoxP, alongside a rise in pGOI-specific content (Table 2). The superior performance likely reflects complete Cre/LoxP recombination and the intrinsic stability of circular products, which are less susceptible to enzymatic degradation and aberrant processing. Collectively, the near-elimination of backbone contamination—while preserving or improving titer—marks a substantive advance for high-purity rAAV manufacturing.

Although TelN-, Cre-, and related strategies reduced plasmid-backbone carryover, applying TelN or Cre to the pRepCap plasmid markedly depressed vector-genome titers and modestly increased residual RepCap sequences in rAAV particles (Figure 2, Figure S5, Table 2). This indicates that cleavage or recombination within—or proximal to—the Rep/Cap expression cassette impairs overall production efficiency. In our pRepCap design, the P5 promoter spans the cap/rep interval (5′ P5 downstream of cap, 3′ P5 upstream of rep), a layout intended for regulatory control and to limit unintended fragment packaging. Nonetheless, the P5 promoter’s proximity to cap and its RBE-TRS–like features likely promote erroneous encapsidation of RepCap and adjacent plasmid sequences. Given Rep helicase/endonuclease activity, plasmid backbones harboring RBE-like motifs near P5 may be processed analogously to ITRs, triggering off-target initiation events—consistent with reports of P5-linked packaging of non-target DNA (Brimble et al., 2022). The concurrent titer loss further suggests that Rep expression is highly sensitive to plasmid architecture. Accordingly, re-engineering Rep/Cap control—e.g., substituting synthetic or heterologous promoters devoid of RBE homology—may reduce contamination risk while preserving rAAV production efficiency. Beyond plasmid-derived contaminants, hcDNA remains a significant challenge in rAAV manufacturing. Such carryover likely arises from non-specific (random) packaging and/or the presence of cryptic RBE-like elements within the host genome, which may engage the Rep protein and cacilitate unintended encapsidation. While our study primarily focused on strategies to minimize plasmid DNA impurities,, the reduction of hcDNA data suggest that suppressing apoptosis during production and incorporating nuclease polishing steps during purification can markedly reduce host-DNA carryover. This supports a model in which apoptosis-induced genomic fragmentation generates substrates for inadvertent packaging. Future efforts should therefore on refining production conditions to minimize cell death and optimize controlled DNA degradation step, thereby further improving rAAV product purity. Our findings have several important implications for improving rAAV manufacturing processes: (1) Plasmid design optimization. Strategically positioning recombination sites (loxP, TelROL) and carefully selecting promoter elements can materially influence both vector purity and yield. Given the pivotal role of the P5 promoter in Rep/Cap carryover, replacing it with alternative or synthetic promoters lacking RBE-like motifs may better balance contamination reduction with titer preservation. (2) Process integration. The Cre/LoxP approach can be seamlessly incorporated into existing workflows—either *via* helper plasmids encoding Cre or Cre-expressing stable producer cell lines—providing a straightforward implementation path that can nearly eliminate backbone contaminants. (3) Scalability. Generating minicircular DNA *in process* during rAAV production obviates extra purification steps required by alternatives (e.g., minicircle or dbDNA workflows), thereby simplifying operations and improving scalability. (4) Regulatory considerations. The Cre/LoxP-driven reduction of plasmid-backbone carryover directly addresses a core CMC concern for clinical-grade vectors, potentially streamlining regulatory review and approval. Despite the advances reported here, several limitations persist and highlight opportunities for further investigation: (1) Vector serotype specificity. Our evaluation centered on AAV9 capsids; the performance of these strategies may differ across other AAV serotypes and warrants validation. (2) Production system dependence.

These approaches were optimized for transient transfection in HEK293-derived cells; their translation to other platforms (e.g., baculovirus/Sf9 or stable producer lines) requires further evaluation. (3) Host-cell DNA reduction. The mechanisms underlying host-genome encapsidation—stochastic capture versus RBE/TRS-mediated events—remain unresolved. Applying single-cell sequencing and chromatin immunoprecipitation (ChIP) to map Rep–host DNA interactions could clarify these pathways and guide targeted countermeasures. (4) Promoter engineering. Designing alternative promoters to drive Rep/Cap expression—explicitly devoid of RBE-like motifs—could further suppress DNA carryover while preserving robust vector productivity. (5) Combination strategies. Integrating complementary DNA-minimization tactics—e.g., pairing Cre/LoxP with nuclease polishing—may further elevate vector purity.

In conclusion, our systematic evaluation of DNA-minimization strategies identifies Cre/LoxP–mediated minicircle formation as a superior approach for substantially reducing plasmid-backbone contamination in rAAV preparations. Coupled with measures to curb host-cell DNA carryover, we present a robust bioprocess toolkit that enhances the safety and purity of rAAV therapeutics and addresses a central manufacturing challenge for clinical applications. Beyond practical gains, these findings deepen mechanistic insight into mispackaging pathways and offer immediately implementable solutions compatible with current production workflows.

## Materials and Methods

### Plasmids

All plasmids were generated using standard molecular biology methods, including gene synthesis, PCR, and Gibson assembly. Constructs were verified by diagnostic restriction enzyme digestion and Sanger sequencing.

#### Gene-of-interest (GOI) vectors

The rAAV genome plasmid pGOI-eGFP comprises the CAG promoter, eGFP reporter, WPRE, and SV40 late polyadenylation signal, flanked by AAV2 ITRs. For pGOI-LucGFP, the firefly luciferase coding sequence followed by the porcine teschovirus 2A (P2A) peptide was synthesized and inserted upstream of eGFP. pGOI-mChe was produced by replacing eGFP in pGOI-eGFP with mCherry.

To enable backbone-segregation strategies, sequence elements were appended to both sides of the ITR-flanked cassette in pGOI-LucGFP:

- TelROL sites upstream of the 5′ ITR and downstream of the 3′ ITR to yield pGOI-TelROL (sequence: *tatcagcacacaattgcccattatacgcgcgtataatggactattgtgtgctgata*).
- I-SceI recognition sites to generate pGOI-IRS (site: *tagggataacagggtaat*).
- loxP sites to generate pGOI-loxP (site: *ataacttcgtatagcatacattatacgaagttat*).

For CRISPR experiments, two custom Tgfp target sites targeting GFP (*ggccacaagttcagcgtgtccgg*) were inserted immediately 5′ of the 5′ ITR and 3′ of the 3′ ITR in pGOI-mChe; an sgRNA against Tgfp was also integrated into the plasmid backbone to create pGOI-Tgfp.

#### AAVS1 targeting vector and expression cassettes

pAAVS1 contains the AAVS1 left homology arm, puromycin resistance, HS4 insulator, CBh promoter, eGFP, bGH polyA, and the right arm. Coding sequences for TelN, I-SceI, SpCas9, and Cre were synthesized and substituted for eGFP in pAAVS1 to generate pTelN, pI-SceI, pCas9, and pCBhpro.Cre, respectively. The CBh promoter in pCBhpro.Cre was further replaced with EFS, hPGK, or UBC to yield pEFSpro.Cre, phPGKpro.Cre, and pUBCpro.Cre.

#### Packaging plasmids

The rAAV packaging plasmid pRepCap encodes AAV2 rep and AAV9 cap, with a P5 promoter positioned downstream of the AAV9 cap gene. For selected experiments, two TelROL sites were inserted upstream of rep and downstream of P5 to create pRepCap-TelROL; analogously, two loxP sites were inserted at the same positions to create pRepCap-loxP.

#### Helper plasmids

The helper plasmid pHelper carries a minimal adenoviral genome segment (E2A, E4, VA RNA) required for AAV replication and packaging. Expression cassettes driven by CBh for TelROL, TelN, or SpCas9 were cloned into pHelper to generate pHelper-TelROL, pHelper-TelN, and pHelper-Cas9. Likewise, Cre expression cassettes under EFS, hPGK, or UBC control were inserted to produce pHelper-EFSpro.Cre, pHelper-hPGKpro.Cre, and pHelper-UBCpro.Cre.

#### sgRNA plasmids

An sgRNA expression vector was assembled by inserting a U6-driven sgRNA cassette into the pUC19 backbone; the Tgfp protospacer (*ggccacaagttcagcgtgtc*) was cloned downstream of U6. To construct pX330-sgAAVS1, the AAVS1 target sequence (*taaggaatctgcctaacagg*) was inserted downstream of the U6 promoter in pX330.

### *In vitro* plasmid digestion/recombination

For *in vitro* separation of plasmid backbones, pGOI-TelROL was treated with TelN protelomerase, pGOI-IRS was digested with the I-SceI meganuclease, and pGOI-loxP was incubated with Cre recombinase. The resulting digested/recombined plasmids were then purified using the E.Z.N.A. Cycle Pure Kit (Omega) prior to rAAV packaging.

### Cell culture and cell line establishment

HEK293T cells were maintained in DMEM (Gibco) supplemented with 10% fetal bovine serum (Gibco) and 1% penicillin–streptomycin (Gibco) at 37 °C in a humidified incubator with 5% CO₂. To generate the KI-I-SceI, KI-Cas9, and KI-Cre cell lines, HEK293T cells were co-transfected with pI-SceI, pCas9, or pCBhpro.Cre together with pX330-sgAASV1 using Lipofectamine 3000 (Invitrogen). Seventy-two hours post-transfection, cells were subjected to puromycin selection for 7–10 days. Puromycin-resistant single-cell clones were then isolated by serial dilution into 96-well plates.

### rAAV vector production

Recombinant AAV (rAAV) particles were generated by co-transfecting HEK293T or KI cells with either a three-plasmid system (pGOI, pRepCap, and pHelper) or a four-plasmid system (pGOI, pRepCap, pHelper, plus an additional accessory plasmid), using PEIpro (Polyplus) as the transfection reagent. For empty rAAV production, only pRepCap and pHelper were co-transfected into HEK293T cells. At 72 h post-transfection, cells were harvested and lysed; crude rAAV preparations were treated with benzonase nuclease to remove residual nucleic acids. Viral particles were then purified by double cesium chloride (CsCl) gradient ultracentrifugation followed by dialysis.

### Titer and hcDNA determination

Capsid titers were quantified by ELISA using an AAV9 capsid–specific antibody. Genome and plasmid-backbone titers were determined by qPCR with WPRE-specific primers (forward, 5′-tgcttcccgtatggctttca-3′; reverse, 5′-acggaattgtcagtgcccaa-3′), ITR-specific primers (forward, 5′- ggaacccctagtgatggagtt-3′; reverse, 5′-cggcctcagtgagcga-3′) and ori-specific primers (forward, 5′-gcgcgtaatctgctgcttg-3′; reverse, 5′-ctacggctacactagaagaacagta-3′), respectively. Residual DNA derived from individual plasmid regions in rAAV preparations was assessed by qPCR using the following primer pairs: Rep (forward, 5′-acgggaactcaacgaccttc-3′; reverse, 5′-ttgcccaccggaaaaagtct-3′), Cap (forward, 5′-ttccactgccacttctcacc-3′; reverse, 5′-ttggcgatggtcttgactcc-3′), E2A (forward, 5′-gagatcagatccgcgtccag-3′; reverse, 5′-ggtgcgagtgcaactcaaag-3′), E4 (forward, 5′-gcatggttgaaggtgctgga-3′; reverse, 5′-ccaaaaggcaaactgccctc-3′), and VARNA (forward, 5′-aatttgcaagcggggtcttg-3′; reverse, 5′-tttccaagggttgagtcgca-3′). The residual hcDNA was measured using a HEK293 residual DNA detection kit.

### DNA NGS sequencing and analysis

Viral DNA was extracted from purified rAAV particles using the HiPure Viral DNA Kit (Magen) following the manufacturer’s instructions. Purified DNA was submitted to Novogene for next-generation sequencing on an Illumina HiSeq platform. Raw reads were processed and aligned to the corresponding plasmid sequences and to the human reference genome for downstream analysis.

### Gene expression analysis

Total RNA was isolated from KI-I-SceI, KI-Cas9, and KI-Cre cell lines using TRIzol reagent (Invitrogen) and reverse-transcribed to first-strand cDNA with the HiScript® III 1st Strand cDNA Synthesis Kit (+gDNA wiper) (Vazyme), following the manufacturers’ instructions. Quantitative RT-PCR (qPCR) was performed with the AceQ Universal SYBR qPCR Master Mix (Vazyme) using the following primer pairs: I-SceI (forward, 5′-ggcggaaagtgggactacaa-3′; reverse, 5′-ctggaacttgttgcgcagtc-3′), SpCas9 (forward, 5′-accatcgaccggaagaggta-3′; reverse, 5′-cgatccgtgtctcgtacagg-3′), Cre (forward, 5′-agaggtatctcgcctggtc-3′; reverse, 5′-cacatttgtccaacctcc-3′), and GAPDH (forward, 5′-ctgggctacactgagcacc-3′; reverse, 5′-aagtggtcgttgagggcaatg-3′). Relative transcript levels were calculated by the 2^−ΔCt method with GAPDH as the internal control.

### Protein expression analysis

To evaluate the expression of I-SceI meganuclease, SpCas9, and Cre recombinase, total cellular proteins were extracted and resolved by SDS–PAGE, followed by transfer to nitrocellulose membranes. Membranes were probed with the appropriate primary antibodies (anti-FLAG or anti-Cas9, as applicable) and HRP-conjugated secondary antibodies. Anti-ACTIN or anti-GAPDH antibodies served as loading controls. Signals were developed with Immobilon Western HRP substrate and imaged on a multi-functional imaging workstation (BLT Biotech).

### Detection of DNA editing activity

Genome-editing activity in KI-Cas9 cell lines was assessed by transiently co-transfecting pGOI-eGFP with either an sgGFP plasmid or a control sgRNA plasmid. At 48 h post-transfection, GFP fluorescence was examined by fluorescence microscopy (Mshot). Genomic DNA was then extracted using the HiPure Universal DNA Kit (Magen). Regions flanking the target site were amplified with GFP-specific primers (forward, 5′-atggtgagcaagggcgagga-3′; reverse, 5′-gtactccagcttgtgcccca-3′). PCR products were denatured and re-annealed to form heteroduplexes, digested with T7 Endonuclease I (Vazyme), and resolved by agarose gel electrophoresis. Gels were imaged on a multifunction imaging workstation (BLT Biotech).

### Investigation of DNA recombination function

Cre-mediated DNA recombination in KI-Cre cell lines was evaluated by transient transfection with either pGOI-LucGFP or pGOI-loxP reporter plasmids. In parallel, HEK293T cells co-transfected with pCBhpro.Cre and either reporter served as controls. At 48 h post-transfection, genomic DNA was isolated using the HiPure Universal DNA Kit (Magen). LoxP-flanking regions were amplified by PCR with the following primer pairs: 5′-ggacaggtatccggtaagcg-3′ / pGOI-R1 (5′-gaggtgccgtaaagcactaa-3′) and pGOI-F2 (5′-ttgggcactgacaattccgt-3′) / pGOI-R2 (5′-agatggggagagtgaagcagaacgt-3′). PCR products were resolved by agarose gel electrophoresis and imaged on a multifunction imaging workstation (BLT Biotech).

## Data availability statement

All data supporting the findings of this study are available within the article and its supplemental information.

## Acknowledgments

We would like to thank many scientists in PackGene Biotech Inc for their technical assistance and contributions.

## Author contributions

J.H designed the study under the guidance of Y.B. and H.L.. J.H., H.C. and Z.D. performed the experiments. X.T. conducted NGS data analysis. J.H. analyzed and prepared the manuscript. J.H., Y.B. and H.L. reviewed the manuscript.

## Supplemental materials

**Figure S1.**
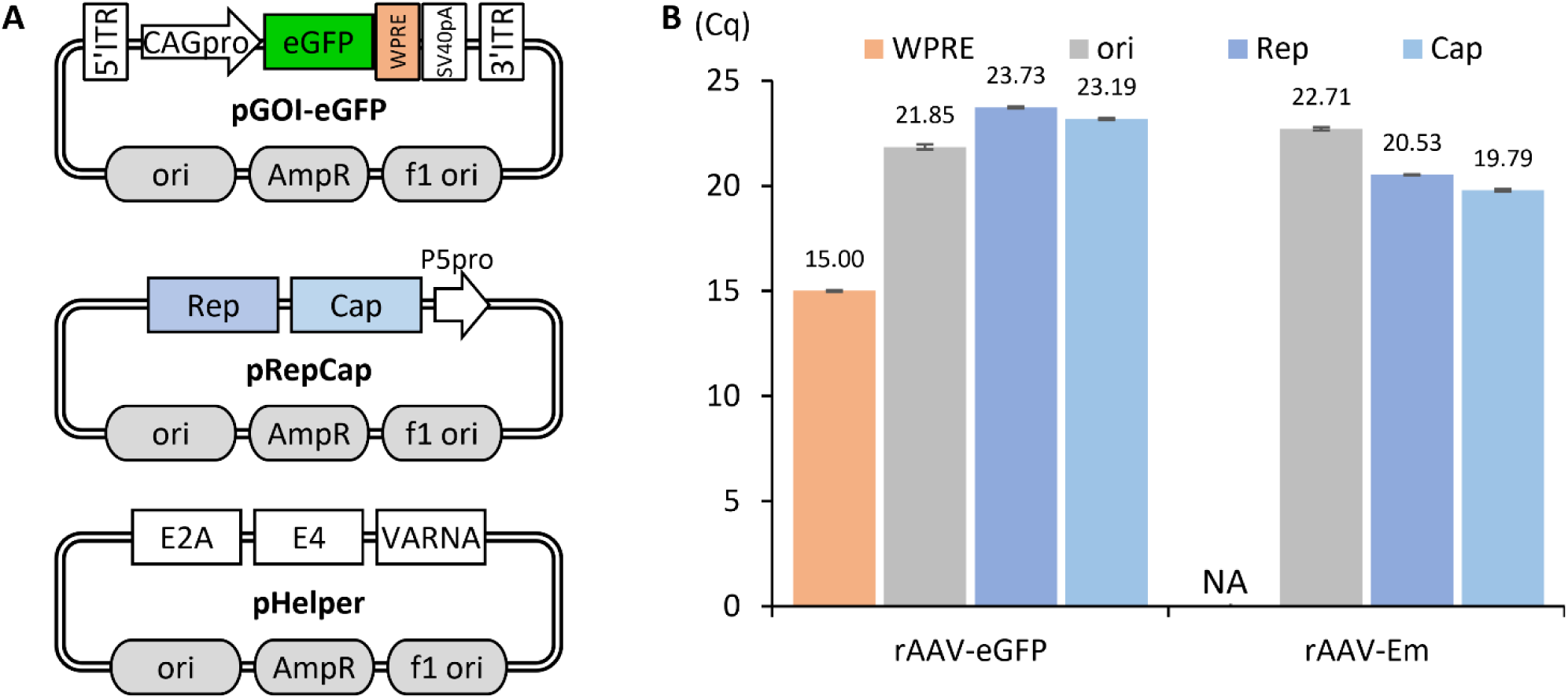
Quantification of residual plasmid DNA in rAAV by qPCR. (A) Schematic maps of the pGOI-eGFP, pRepCap, and pHelper plasmids. (B) Quantification cycle (Cq) values for WPRE, ori, Rep, and Cap targets in purified rAAV samples. rAAV-eGFP: co-transfection of pGOI-eGFP, pRepCap, and pHelper in HEK293T cells. rAAV-Em: co-transfection of pRepCap and pHelper in HEK293T cells. Cq, quantification cycle in qPCR (lower Cq indicates a higher template amount). NA, data not available.

**Figure S2.**
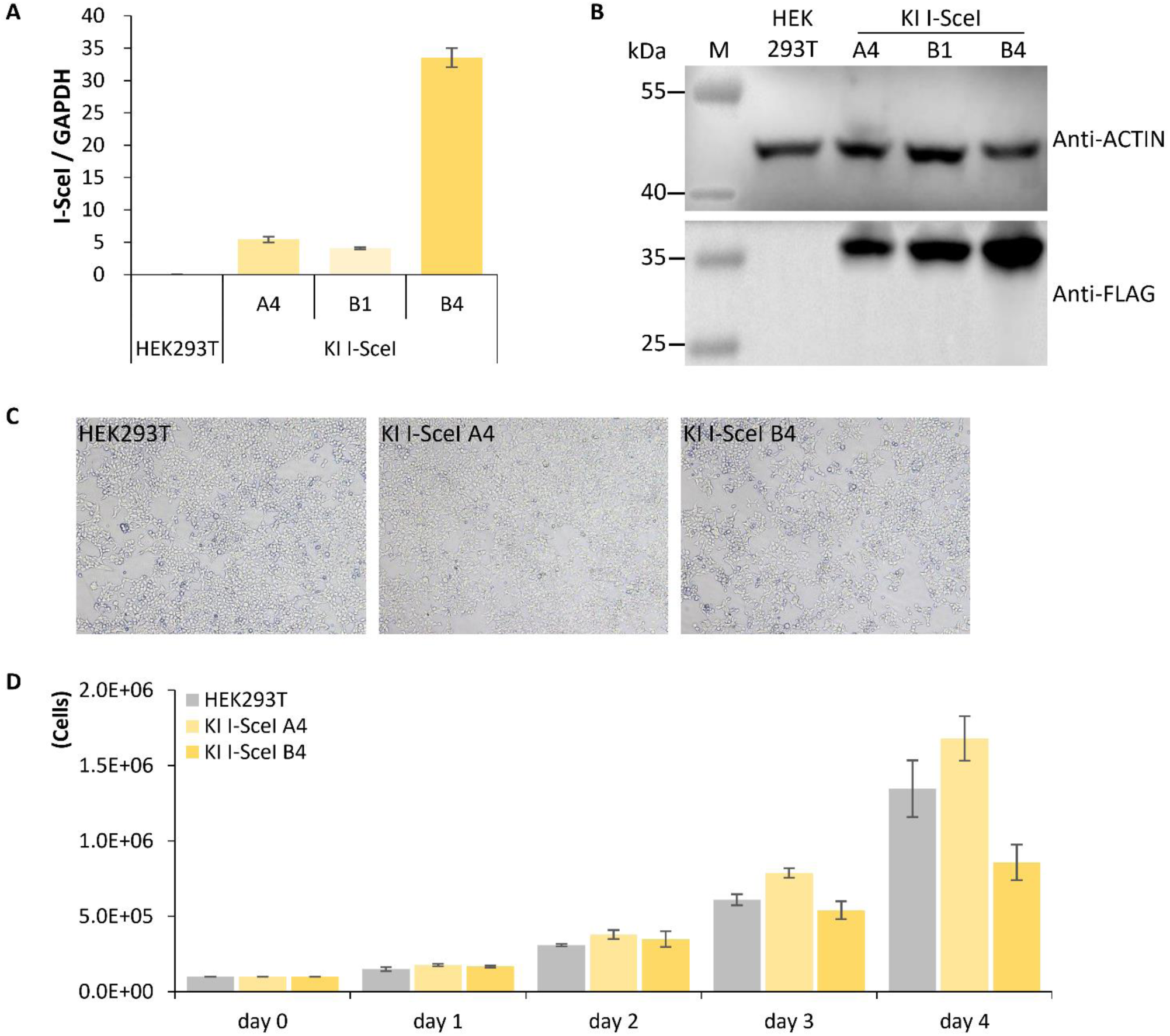
Detailed characterization of I-SceI Knock-in (KI I-SceI) cell lines. (A, B) Expression levels of I-SceI mRNA (A) and protein (B) in the KI I-SceI cell lines were quantified by qPCR and western blot analysis, respectively. (C, D) The cell morphology (C) and growth kinetics (D) of the KI I-SceI cell lines were evaluated.

**Figure S3.**
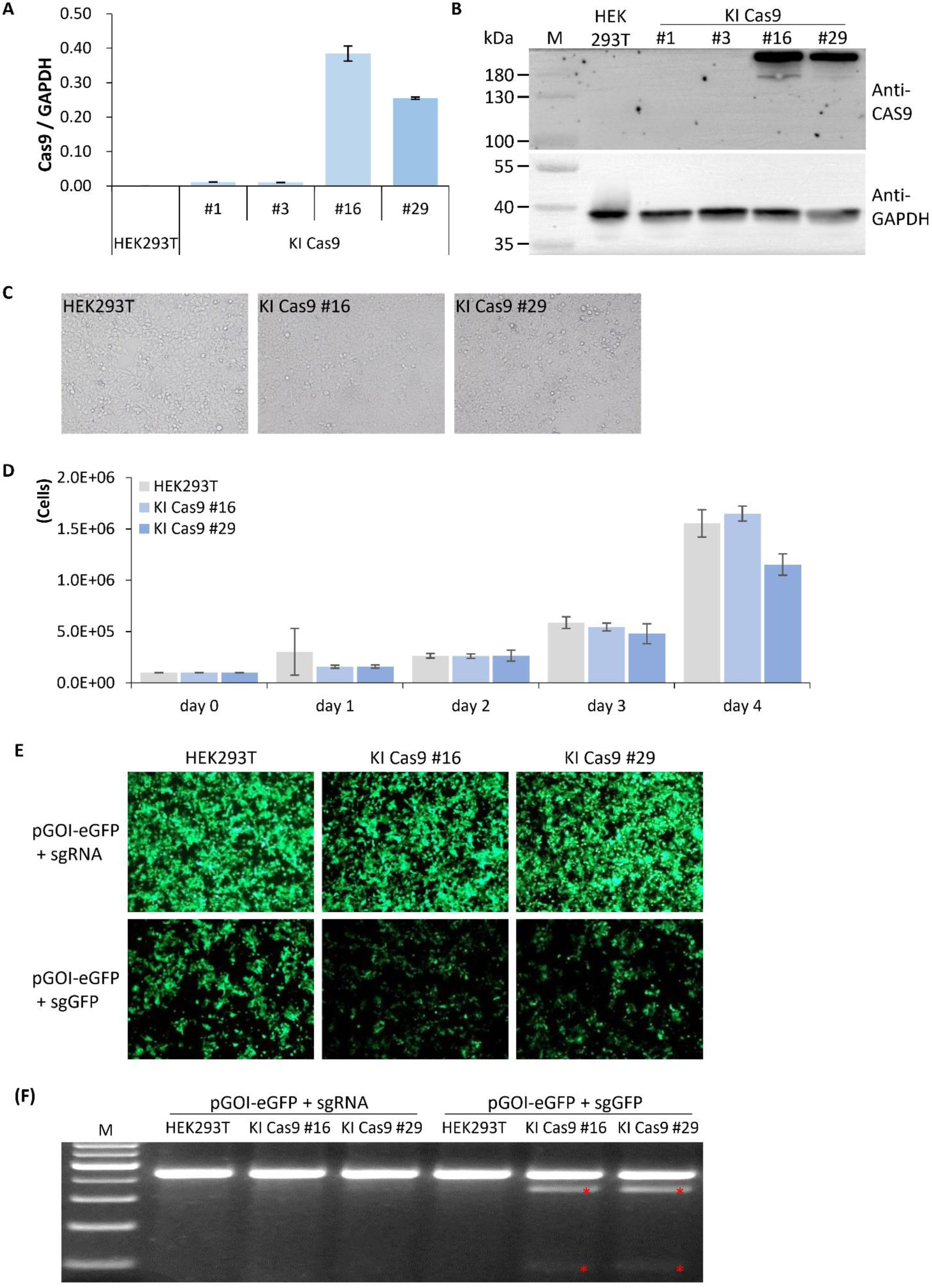
Generation and characterization of Cas9 Knock-in (KI-Cas9) cell lines. (A, B) Expression levels of Cas9 mRNA (A) and protein (B) in the KI Cas9 cell lines were quantified by qPCR and western blot analysis, respectively. (C, D) The cell morphology (C) and proliferation rate (D) of the KI Cas9 cell lines were assessed. (E, F) Genome-editing activity was evaluated by co-transfecting pGOI-eGFP with an sgGFP plasmid. GFP signals were visualized by fluorescence microscopy (E), and cleavage efficiency was determined by T7 endonuclease I (T7E1) assay (F); red asterisks indicate cleaved fragments.

**Figure S4.**
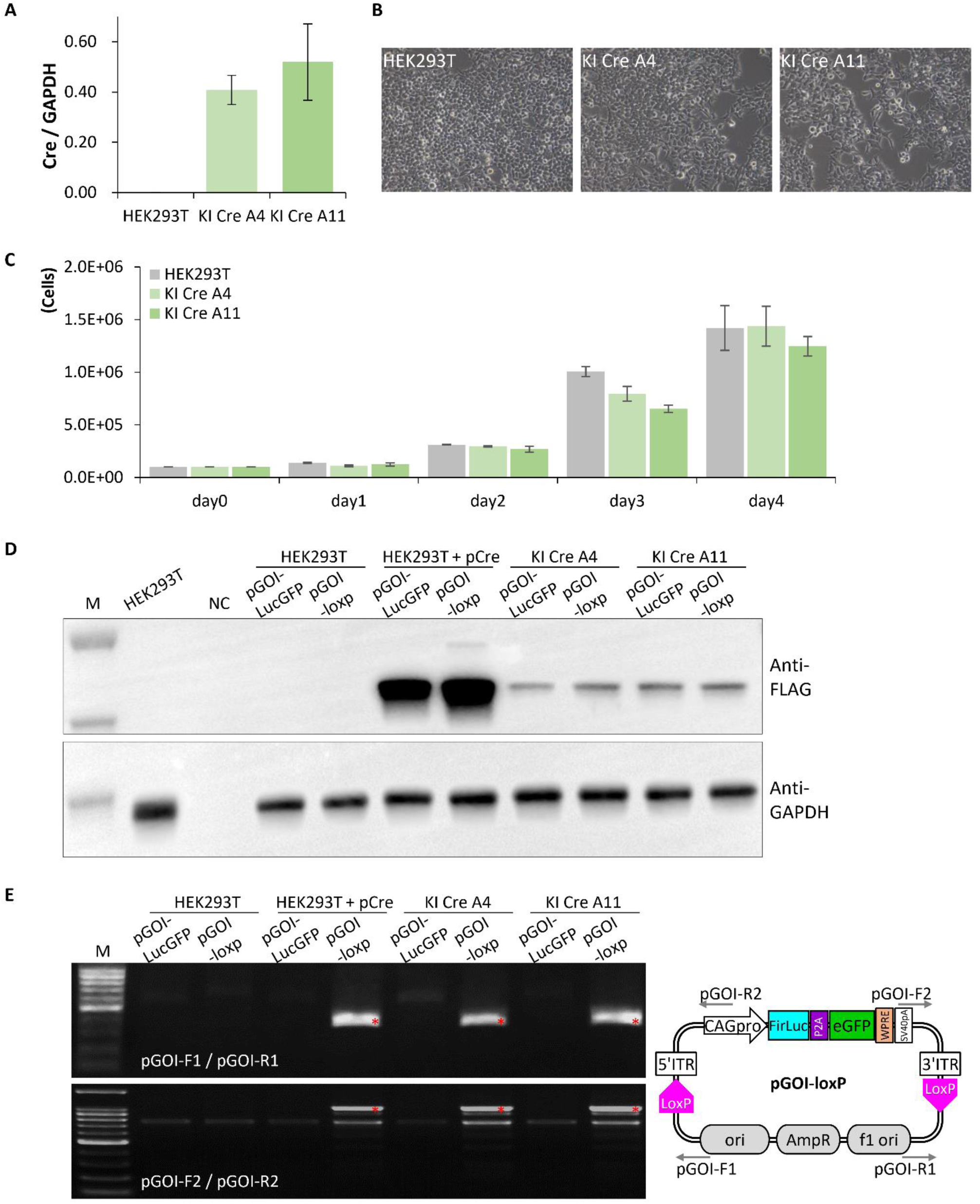
Establishment and characterization of Cre Knock-in (KI-Cre) cell lines. (A) Cre transcript levels in KI-Cre cell lines quantified by qPCR. (B) Representative cell morphology of KI-Cre lines. (C) Proliferation rates of KI-Cre lines. (D, E) Cre-mediated recombination was assessed by transient transfection of pLucGFP-loxP; co-transfection of pCre and pLucGFP served as a positive control. Cre protein expression was examined by Western blot (D), and recombination events were detected by PCR (E). red asterisks indicate recombined circular DNA.

**Figure S5.**
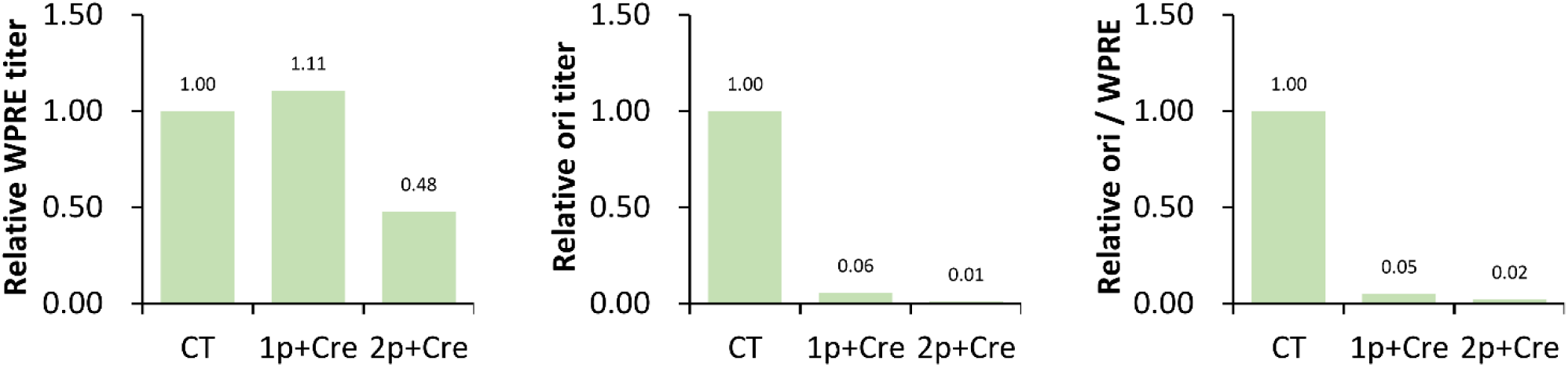
Assessment of DNA impurities in rAAV produced *via* the Cre/LoxP system for NGS analysis. rAAV was generated using pGOI-loxP, pRepCap-loxP, and pHelper-Cre. Experimental conditions were as follows: CT, co-transfection of pGOI-loxP/pRepCap/pHelper; 1p+Cre, co-transfection of pGOI-loxP/pRepCap/pHelper-Cre; 2p+Cre, co-transfection of pGOI-loxP/pRepCap-loxP/pHelper-Cre. The genome titer (WPRE) and plasmid-backbone titer (ori) were quantified by qPCR. Relative WPRE, relative ori, and the ori/WPRE ratio were calculated against the corresponding control rAAV samples.

